# miR-495-3p inhibition rescues mTORC1 hyperactivation-driven autistic-like behaviors in mice

**DOI:** 10.64898/2026.03.02.708955

**Authors:** Brunno R. Levone, Nik Schneider, Paulina Deltuvaite, Pierre-Luc Germain, Gerhard Schratt

## Abstract

Defective social behavior and cognitive functions are hallmarks of Autism spectrum disorders (ASD). The molecular mechanisms keeping social behavior in a physiological range are largely unknown. We recently found that conditional knockout (cKO) of tmiRNA cluster miR-379-410 in mouse hippocampal neurons leads to hypersocial behavior. Therefore, inhibiting miR-379-410 members might represent a strategy to promote sociability in ASD. As an ASD model, we chose knockdown (KD) of the ASD risk gene Tsc1, a key negative regulator of mTORC1. Acute Tsc1 knockdown (KD) in hippocampal neurons was sufficient to induce hyposociability and memory deficits in adult wild-type mice. In contrast, Tsc1 KD had no effect on sociability in miR-379-410 cKO mice, indicating a requirement of miR-379-410. Furthermore, Tsc1 KD led to upregulation of tmiR-495-3p, and inhibition of this miRNA by antisense oligonucleotides was sufficient to prevent hyposociability and memory impairments. Our findings suggest that miR-495-3p is a key downstream effector of the Tsc1/mTORC1 pathway in sociability, and that targeting miR-495-3p represents a therapeutic avenue for restoring social and cognitive impairments in ASD without affecting mTORC1 homeostasis.

## 1. Introduction

Autism spectrum disorder (ASD) and hypersocial syndromes (such as Williams-Beuren syndrome-WBS) are situated at opposite ends of a spectrum of social behavioral traits. In ASD, core symptoms include impaired sociability, restricted interests, and repetitive behaviors. Conversely, hypersocial phenotypes are characterized by exaggerated sociability and diminished social inhibition. Altered development and plasticity of neural circuits involved in the control of social behavior underlies social dysfunction, likely as a result of aberrations in gene regulatory networks in opposite directions. Emerging data suggest that microRNAs (miRNAs) are dysregulated in samples from both patients with ASD (Wu, Parikshak et al. 2016) and with WBS (Kimura, Swarup et al. 2019), but whether they contribute to the social phenotypes observed in these diseases is not yet clear.

MiRNAs, small non-coding ribonucleic acids (RNAs), play key roles in post-transcriptional gene regulation by binding to target messenger RNAs (mRNAs) and directing them to degradation or translational repression. Thus, miRNAs are known as fine-tuners of gene networks, influencing diverse biological processes including neural development, synaptic plasticity, and behavioral response (Bartel 2018). Indeed, miRNAs serve as dynamically responsive molecular switches linking environmental input and genetic content with long-term cellular and circuit changes (Flynt and Lai 2008). The miR-379-410 cluster, located at the imprinted *Dlk1*-*Dio3* locus and expressed exclusively from the maternal allele, is a miRNA cluster with several known roles in neurons. Our laboratory has recently described that the knockout of this cluster in mice results in hypersocial and anxiogenic phenotypes, along with changes in excitatory synaptic transmission (Lackinger, Sungur et al. 2019). More recently, we showed that the conditional knockout of this cluster in excitatory forebrain neurons (*Emx1*-expressing) drives a similar hypersocial behavior but only in adult male mice. Hypersocial behavior is accompanied by increased excitatory synaptic transmission and enhanced expression of acto-myosin genes in the hippocampal CA1 pyramidal neurons (Narayanan, Levone et al. 2024). On the other hand, overexpression of specific miR-379-410 members induces hyposocial behavior. These findings indicate that the miR-379-410 cluster acts as a brake on sociability, integrating synaptic structure and neural circuit function into behavioral outcomes. However, upstream signaling pathways controlling miR-379-410 expression in the brain in the context of social behavior remain largely unknown.

A growing body of evidence links the mammalian target of rapamycin complex 1 (mTORC1) pathway to the control of social behavior. mTORC1 is at the nexus of nutrient sensing, growth-factor signaling, and synaptic plasticity, and is critically involved in shaping neuronal connectivity and function. An increased mTORC1 signaling is usually described in ASD patients and mouse models (Drehmer, Santos-Terra et al. 2024), and the hyperactive mTORC1 phenotypes generally involve exaggerated protein synthesis, mitochondrial dysfunction, and altered dendritic spine morphology (Nicolini, Ahn et al. 2015, Magdalon, Sanchez-Sanchez et al. 2017, Pagani, Barsotti et al. 2021). For instance, Tuberous Sclerosis, a monogenic disorder caused by mutations in the *Tsc1* or *Tsc2* genes, leads to constitutive mTORC1 activation, and is frequently associated with comorbidities such as epilepsy and autistic behavior. In *Tsc1*-and *Tsc2*-mutant mice, treatment with the mTORC inhibitor rapamycin rescues social behavior deficits, implicating mTORC1 as a targetable hub for social regulation (Sato, Kasai et al. 2012).

Genetic mouse models of *Tsc1* provide a powerful system for investigating how mTORC1 hyperactivation influences neural function and behavior. For example, *Tsc1* haploinsufficiency in parvalbumin (PV) interneurons impairs social approach and social memory, while complete *Tsc1* deletion in these cells additionally leads to an anxiolytic effect (Amegandjin, Choudhury et al. 2021). Loss of *Tsc1* in cerebellar Purkinje cells induces autistic-like behaviors in mice, a phenotype that can be rescued by rapamycin treatment (Tsai, Hull et al. 2012). Similarly, *Tsc1* haploinsufficiency in medial ganglionic eminence-derived inhibitory neurons (Nkx2.1-positive) reduces synaptic inhibition onto pyramidal cells, leading to deficits in contextual fear discrimination and spatial working memory (Haji, Riebe et al. 2020). In the hippocampus, *Tsc1* is required to maintain excitatory synaptic strength, and its loss abolishes metabotropic glutamate receptor-dependent long-term depression, a key form of synaptic plasticity (Bateup, Takasaki et al. 2011). Moreover, *Tsc1* loss-of-function in forebrain excitatory neurons using Camk2a-Cre causes frequent seizures and has been proposed as a mouse model of epilepsy (Koene, van Grondelle et al. 2019), However, the effects on social behavior caused by hippocampal-specific *Tsc1* loss-of-function have not been addressed.

Although *Tsc1* has been manipulated in many neuronal populations, relatively few studies have explored the molecular mechanisms leading to social impairments downstream of *Tsc1*. Interestingly, in the mouse liver, loss of *Tsc1* leads to mTORC1 activation and the upregulation of several miRNAs, with particular enrichment of miRNAs encoded within the imprinted *Dlk1*-*Dio3* locus, including the miR-379-410 cluster (Liko, Rzepiela et al. 2020). Based on these findings, we hypothesized that *Tsc1* knockdown may similarly drive upregulation of the miR-379-410 cluster in mouse hippocampal excitatory neurons, which in turn would cause gene expression changes leading to hyposociability. If true, directly targeting miR-379-410 members (e.g., using antisense oligonucleotides (ASOs)) could represent a novel therapeutic strategy to mitigate autism-related phenotypes associated with *Tsc1* loss, such as hyposociability, cognitive impairments and potentially epileptic seizures.

## 2. Results

### 2.1. Acute *Tsc1* knockdown in excitatory neurons of the hippocampus impairs social behavior and recognition memory in mice

The miR-379-410 acts as a brake for sociability in the adult mouse hippocampus (Lackinger, Sungur et al. 2019, Narayanan, Levone et al. 2024). To investigate a potential link between *Tsc1* and miR-379-410 in the hippocampus, we first decided to re-visit the role of hippocampal *Tsc1* in social behavior. Conditional *Tsc1* knockout (cKO) in forebrain excitatory neurons (Camk-Cre deletion) leads to a strong hyperexcitability and the occurrence of hippocampal seizures (Bateup, Johnson et al. 2013), which hampers investigation of a potential contribution of hippocampal excitatory neurons to specific behaviors, such as sociability and memory. To circumvent this problem, we decided to modestly and acutely reduce *Tsc1* expression in adult excitatory hippocampal neurons using shRNA-mediated *Tsc1* silencing. The efficiency and specificity of the shRNA sequence used for *Tsc1* knockdown (KD) was initially validated in Neuro2a cells (see Suppl. Results and Suppl. Figure 1A-J). Subsequently, adult (postnatal week [PNW] 12-15) C57BL/6 wild-type mice underwent stereotaxic surgery for bilateral injection of a recombinant adeno-associated virus (rAAV) expressing the *Tsc1*-Sh or control shRNA (Ctrl-Sh) under the control of the *Camk2a* promoter into the dorsal/intermediate hippocampus. Three weeks post-surgery, mice were subjected to a battery of behavioral tests (timeline in Figure 1A). While *Tsc1*-Sh expression induced a robust hyposocial phenotype in female mice (Figure 1B), it surprisingly did not affect sociability in the reciprocal social interaction test in male mice (Figure 1C). The sex-specific effect of *Tsc1* KD on social behavior was corroborated with an orthogonal test, the 3-chambers test (Figure 1D-E). Interestingly, male mice showed a mild, non-significant trend towards increased social interaction in the reciprocal social interaction test, which may reflect a confounding effect on locomotor activity specifically observed in male pairs (Suppl. Figure 2A-B). Neither female nor male mice showed alterations in social novelty preference in the three-chambers test (Suppl. Figure 2C-D). Interestingly, *Tsc1*-Sh impaired short-term recognition memory in both female and male mice, as assessed using the novel object preference test (Figure 1F-G), suggesting sex-independent effects of *Tsc1* knockdown on cognitive function.

**Figure 1:**
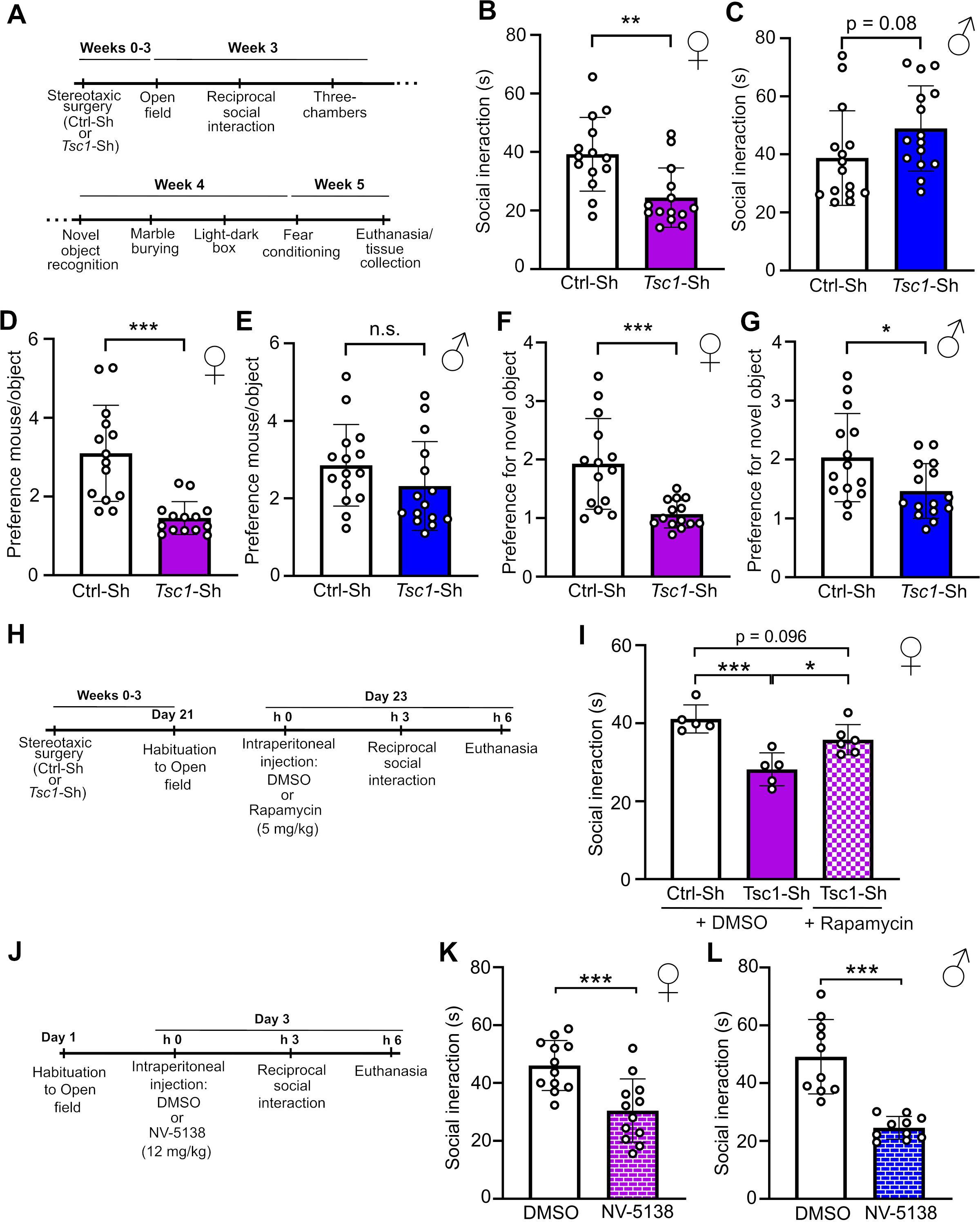
*Tsc1* knockdown in the hippocampus of adult female mice induced hyposocial behavior in an mTORC1-dependent manner. **A.** Timeline of the *Tsc1* knockdown behavioral experiment. Mice underwent stereotaxic surgery for the injection of rAAV-PHP.eB expressing either a Control (Ctrl) or *Tsc1* short hairpin (Sh) into the dorsal/intermediate hippocampus. After 3 weeks, mice were subjected to a battery of behavioral tests. **B-C.** Reciprocal interaction test. While *Tsc1* knockdown reduced social behavior in female mice (B), it led to a trend to increase social behavior in male mice (C). **D-E.** Three-chambers test – sociability. While *Tsc1* knockdown reduced sociability in female mice (D), it did not change it in male mice (D). **F-G.** Novel object recognition. *Tsc1* knockdown decreased the preference for the novel object in both female (F) and male mice (G). **H.** Timeline of the *Tsc1* knockdown with Rapamycin treatment experiment. Female mice underwent stereotaxic surgery for the injection of rAAV-PHP.eB expressing either a Ctrl or *Tsc1* Sh into the dorsal/intermediate hippocampus. After 3 weeks, mice were injected intraperitoneally with either DMSO or Rapamycin (5 mg/kg) diluted in saline. Social behavior was tested in the reciprocal social interaction test 3 h later. **I.** Reciprocal interaction test. The acute (3 h) inhibition of mTORC1 activity with rapamycin rescued *Tsc1* knockdown-induced reductions in social behavior in female mice. **J.** Timeline of the acute NV-5138 (mTORC1 activator) treatment experiment. Mice were injected intraperitoneally with either DMSO or NV-5138 (12 mg/kg) diluted in saline. Social behavior was tested in the reciprocal social interaction test 3 h later. **K-L**. Reciprocal interaction test. The acute (3 h) activation of mTORC1 pathway by intraperitoneal injection of NV-5138 decreased social behavior in both female (K) and male mice (L).

In the open-field test, neither females nor males displayed any shRNA-dependent alterations in locomotor activity or in time spent in the center of the arena (Suppl. Figure 2E-H). Similarly, *Tsc1*-Sh did not affect behavior in the marble-burying test (number of marbles buried; Suppl. Figure 2I-J), in the light-dark box test (number of transitions; Suppl. Figure 2K-L), or in the fear-conditioning test (% freezing; Suppl. Figure 2M-N).

To test whether *Tsc1* knockdown-induced effects in social behavior are indeed dependent on the activation of the mTORC1 pathway, *Tsc1*-Sh-injected female mice were intraperitoneally injected with the mTORC1 inhibitor, rapamycin, 3 h prior to the performance of the reciprocal social interaction test (timeline in Figure 1H). Hyposocial behavior upon *Tsc1* KD was not observed in rapamycin-injected mice (Figure 1I), confirming that *Tsc1* mediates its effect on sociability via mTORC1 inhibition. Moreover, acute mTORC1 activation by intraperitoneal injection of the mTORC1 activator NV-5138 (Sengupta, Giaime et al. 2019) (timeline in Figure 1J) decreased social behavior both in female (Figure 1K) and male mice (Figure 1L), demonstrating that acute hyperactivation of mTORC1 is sufficient to decrease sociability. Neither rapamycin (Suppl. Figure 2O) nor NV-5138 (Suppl. Figure 2P-Q) changed the locomotor activity of the pairs during the reciprocal social interaction test.

Taken together, our results indicate that acute *Tsc1* knockdown in excitatory hippocampal neurons of adult female mice is sufficient to induce hyposocial behavior in an mTORC1-dependent manner.

### 2.2. *Tsc1* knockdown led to an upregulation of miR-495-3p in the hippocampus of mice

After completion of the behavioral assessments, mice were euthanized, and brains were collected for molecular and histological analyses. As visible by unstained brain slices, viral spread (GFP-expressing neurons) reached the CA1 and CA2 regions of the dorsal hippocampus, with very little spread into the CA3, dentate gyrus, and ventral hippocampus (Figure 2A). Because *Tsc1* silencing was restricted to excitatory neurons transduced by the rAAV, only a modest reduction in bulk *Tsc1* expression was expected. Consistently, real-time quantitative PCR (RT-qPCR) revealed only a marginal decrease in *Tsc1* mRNA levels (-16% relative to Ctrl-Sh, Suppl. Figure 3A), while western blot analysis showed an even smaller trend at the protein level (-10% relative to Ctrl-Sh, Suppl. Figure 3B-C). To specifically assess TSC1 expression in transduced neurons, we performed immunohistochemistry and observed that TSC1 immunoreactivity was significantly reduced (about 40%) in rAAV-infected neurons within the hippocampus (Figure 2B-C), confirming effective local silencing of *Tsc1* to levels which are not expected to result in epileptic seizures.

**Figure 2:**
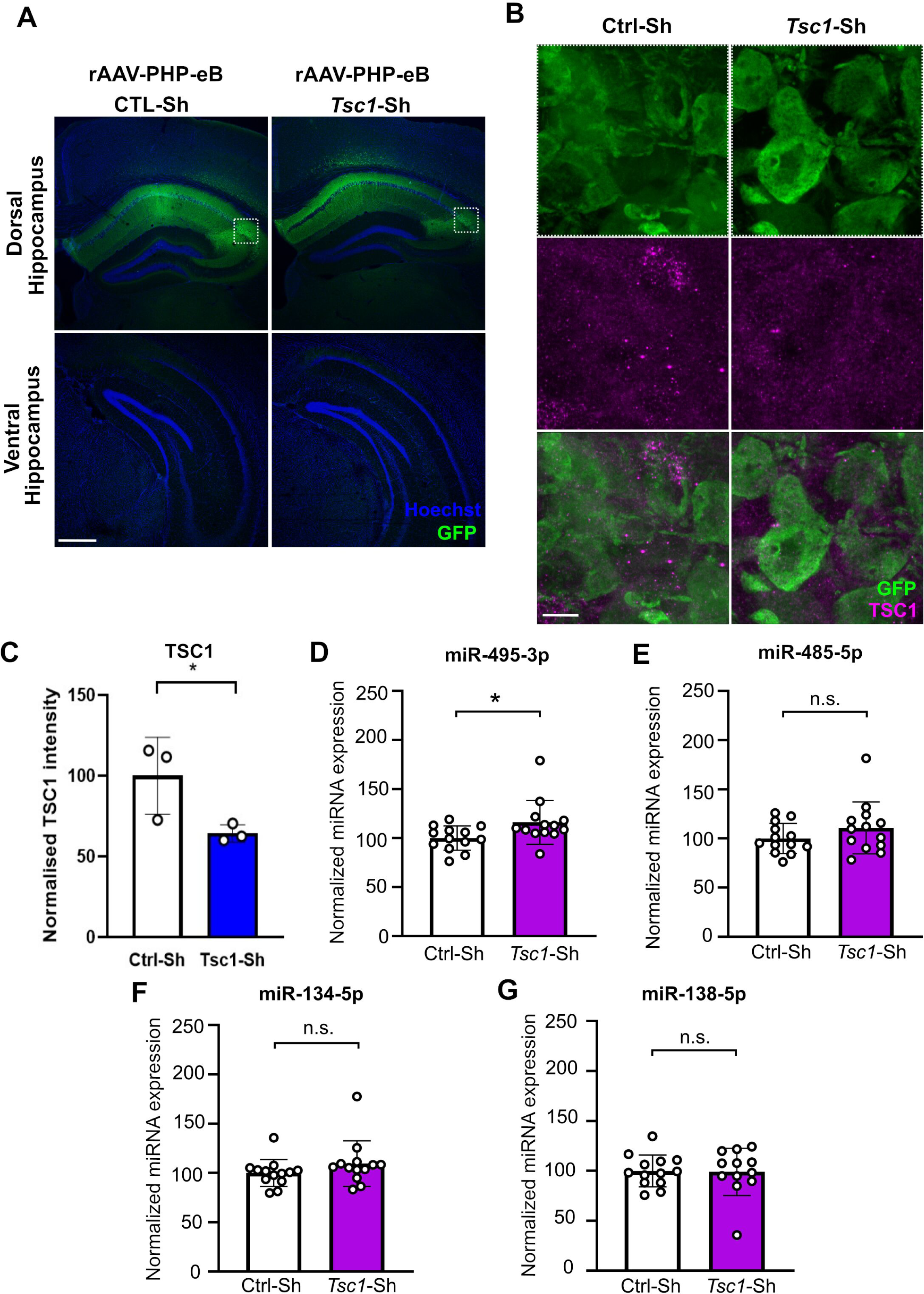
*Tsc1* knockdown induced an increase in miR-495-3p expression in the adult mouse hippocampus. **A**. Representative pictures of the viral expression (rAAV-PHP.eB Ctrl-Sh and *Tsc1*-Sh) in the dorsal (top panels) and ventral (bottom panels) hippocampus. Scale bar: 500 µm. **B-C**. Immunohistochemistry in brain slices of mice injected with either Ctrl-Sh or *Tsc1*-Sh. TSC1 protein was decreased at the viral expression location in the hippocampus of *Tsc1*-Sh-injected mice. Scale bar: 10 µm. **D-F**. While the expression of miR-495-3p (D) was upregulated upon *Tsc1* knockdown, the expression of miR-485-5p (E), and miR-134-5p (F) were unchanged. **G**. The expression of miR-138-5p, used as a negative control, was unchanged upon *Tsc1* knockdown.

In the liver, *Tsc1* cKO leads to a strong upregulation of several members of the miR-379-410 cluster (Liko, Rzepiela et al. 2020). To assess whether *Tsc1* knockdown has similar effect in the adult hippocampus, we performed TaqMan RT-qPCR analyses for specific cluster members. For this analysis, we focused on miR-495-3p, miR-134-5p, and miR-485-5p, since our previous studies (Lackinger, Sungur et al. 2019, Narayanan, Levone et al. 2024) showed that these miRNAs have an important role in the regulation of social behavior and synaptic genes in miR-379-410 KO mice. This analysis showed that miR-495-3p levels were significantly upregulated in the hippocampus of *Tsc1* KD compared to control mice (Figure 2D). In contrast, the expression of miR-485-5p and miR-134-5p were unchanged (Figure 2E-F). We observed no changes in the expression of miR-138-5p, which is not a member of the cluster and was used as a negative control (Figure 2G). Similar results were observed upon *Tsc1* KD in Neuro2A cells (Suppl. Figure 1), although miR-495-3p upregulation was more pronounced and miR-485-5p was also upregulated, likely due to the higher *Tsc1* knockdown efficiency (> 70%) in these cells. Together, these data suggest that the expression of at least one member of the miR-379-410 cluster, miR-495-3p, is induced in neurons upon *Tsc1* knockdown.

### 2.3. *Tsc1* knockdown fails to induce hyposocial behavior in miR379-410 conditional KO mice

Next, we sought to assess the functional involvement of miR-379-410 cluster miRNAs in *Tsc1* knockdown-induced hyposocial behavior. Therefore, we made use of the previously described Emx1-flox miR-379-410 conditional knockout (cKO) mice (Narayanan, Levone et al. 2024), reasoning that shRNA-mediated *Tsc1* knockdown should have no effect on social behavior if the expression of miR-379-410 members were required downstream of *Tsc1*. As controls (CTL), we used mice containing a floxed miR-379-410 allele, but without Cre expression, which display unaltered miR-379-410 expression. We injected the rAAV expressing the *Tsc1*-Sh in CTL and cKO mice at PNW 12 and submitted mice to a battery of behavioral testing three weeks later (timeline in Figure 3A). In female CTL mice, *Tsc1* knockdown in hippocampal excitatory neurons robustly induced hyposocial behavior as assessed by both the social interaction (Figure 3B) and three chambers test (Figure 3C), which is consistent with the results described above (Figure 1B/1D). However, this effect was completely abolished in miR-379-410 cKO mice (Figure 3B-C). Importantly, *Tsc1* knockdown led to reduced novel object discrimination (Figure 3D) in both CTL and miR-379-410 cKO mice, suggesting that the lack of the entire miR-379-410 cluster is specifically required for social behavior but not cognitive performance. In male mice, *Tsc1* knockdown effects were more variable, making it more difficult to draw specific conclusions about the contribution of miR-379-410 regulation in this sex (Suppl. Figure 4A-C). *Tsc1*-Sh did not affect locomotor activity in any of the experimental groups, either during the reciprocal social interaction test (Suppl. Figure 4D-E) or in the open field test (Suppl. Figure 5A-B). It also did not affect anxiety behavior, measured by the time spent in the center of the open field (Suppl. Figure 5C-D), the number of marbles buried (Suppl. Figure 5E-F) and in the light-dark box (Suppl. Figure 5G-H). Taken together, miR-379-410 expression is required for hyposocial behavior, but not impaired memory, caused by *Tsc1* knockdown in hippocampal excitatory neurons of female mice.

**Figure 3:**
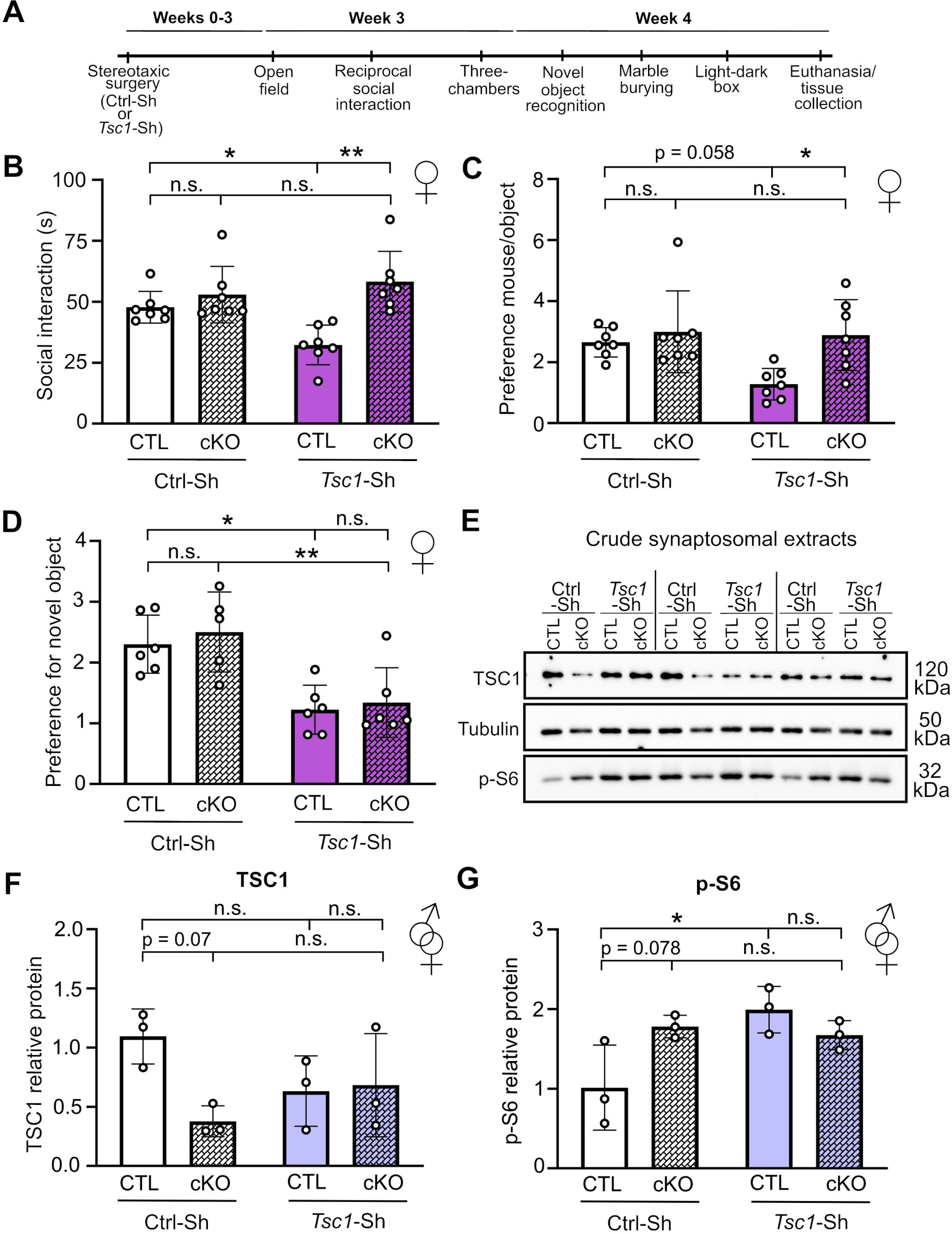
The miR-379-410 cluster is required for hyposociability caused by *Tsc1* knockdown in the hippocampus. **A.** Timeline of the *Tsc1* knockdown behavioral experiment. Mice underwent stereotaxic surgery for the injection of rAAV-PHP.eB expressing Ctrl or *Tsc1* Sh into the dorsal/intermediate hippocampus. After 3 weeks, mice were subjected to a battery of behavioral tests. **B.** Reciprocal interaction test. While *Tsc1* knockdown decreased social behavior in control (CTL) female mice, it did not affect social behavior in mice with a conditional knockout (cKO) of the full miR-379-410 cluster in excitatory neurons of the forebrain. **C.** Three-chambers test – sociability. Similarly to the reciprocal social interaction test, *Tsc1* knockdown only reduced sociability in CTL, but not cKO female mice. **D.** Novel object recognition. *Tsc1* knockdown decreased the preference for the novel object both in CTL and in cKO female mice. **E-G**. Crude synaptosomal extracts show that TSC1 protein is already less expressed in cKO mice, and that *Tsc1* knockdown marginally decreases TSC1 protein in CTL, but not in cKO mice. In agreement, cKO mice have increased phosphorylation of S6, and *Tsc1* knockdown increases phosphorylation of S6 in CTL, but does not further increase it in cKO mice.

Next, we investigated how the observed behavioral changes correlated with the activity of the mTORC1 pathway by measuring TSC1 and phospho-S6 protein levels in crude synaptosomal extracts by Western blotting. Interestingly, while TSC1 knockdown leads to the expected decrease in *Tsc1* and a corresponding increase in phospho-S6 in CTL mice, these changes are not observed in cKO mice (Figure 4E-G). The absence of effect in cKO mice is likely due to lower baseline levels of *Tsc1*, which in turn correspond with elevated baseline p-S6 levels (Figure 4E-G). The reason for the lower *Tsc1* levels upon miR-379-410 knockout are currently unknown, but could involve a compensatory feedback regulation in response to hypersociability.

**Figure 4:**
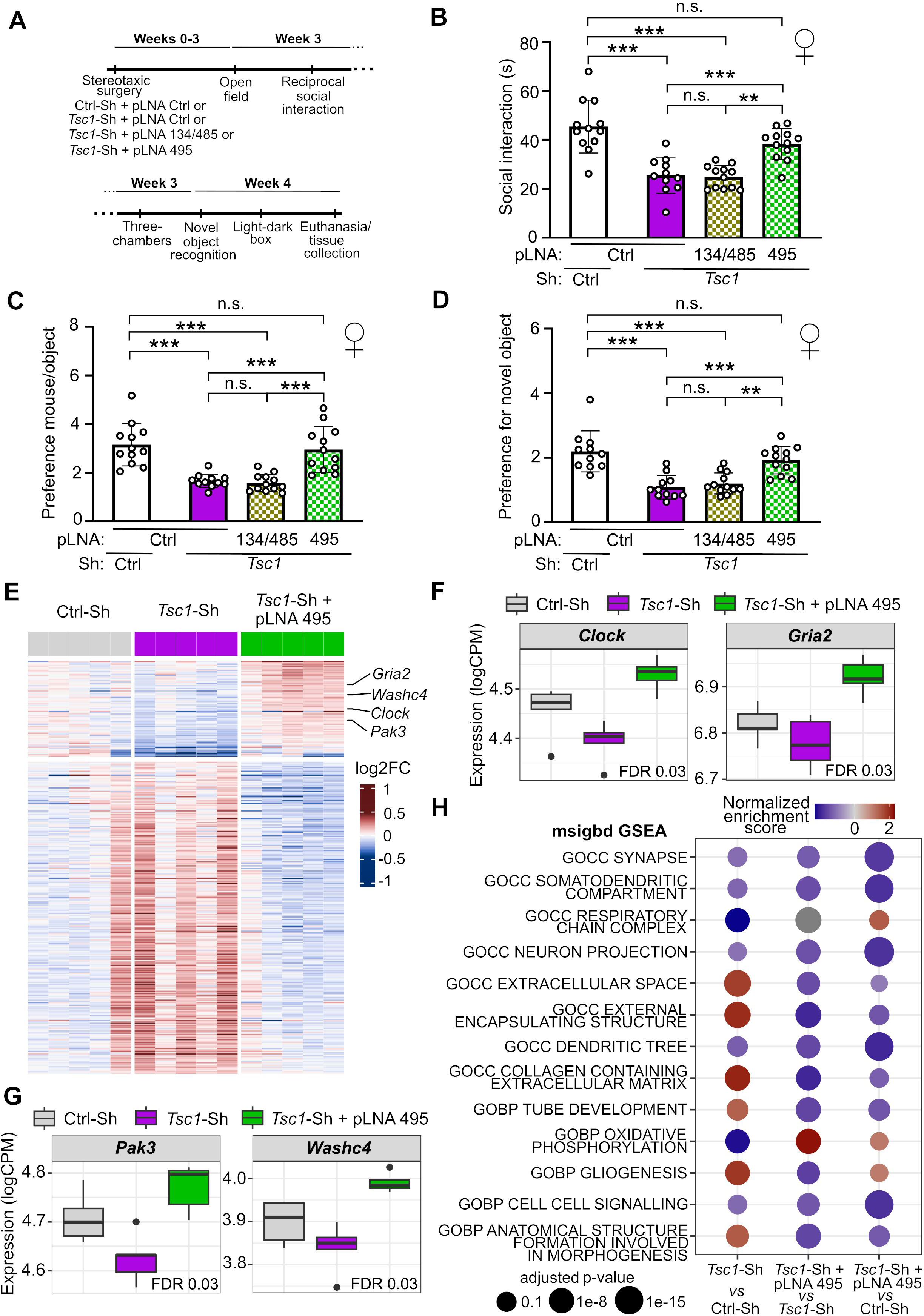
Antisense-oligonucleotide (ASO) mediated inhibition of miR-495-3p rescues hyposociability caused by *Tsc1* knockdown in the hippocampus. **A.** Timeline of the *Tsc1* knockdown behavioral experiment. Mice underwent stereotaxic surgery for the injection of rAAV-PHP.eB expressing Ctrl or *Tsc1* Sh, together with a pLNA (either control-Ctrl, or against miR-495-3p, or against miR-134-5p together with miR-485-5p) into the dorsal/intermediate hippocampus. After 3 weeks, mice were subjected to a battery of behavioral tests. **B.** Reciprocal interaction test. The specific inhibition of miR-495-3p, but not of miR-134-5p/miR-485-5p, prevented *Tsc1* knockdown-induced reductions in social behavior. **C.** Three-chambers test – sociability. Similarly, the specific inhibition of miR-495-3p, but not of miR-134-5p/miR-485-5p, prevented *Tsc1* knockdown-induced reductions in sociability. **D.** Novel object recognition. Similarly, the specific inhibition of miR-495-3p, but not of miR-134-5p/miR-485-5p, prevented *Tsc1* knockdown-induced reductions in the preference for the novel object. **E.** Heatmap of the top dysregulated genes among all groups. There is a clear pattern for the group *Tsc1*-Sh with pLNA against miR-495-3p to be more similar to the Ctrl-Sh than the *Tsc1*-Sh group. **F-G.** *Clock*, *Gria2*, *Pak3*, and *Washc4* genes are some of the examples of the genes downregulated by *Tsc1*-Sh that are rescued by the inhibition of miR-495-3p. The false discovery rate (FDR) values for the comparison *Tsc1*-Sh *vs Tsc1*-Sh with pLNA against miR-495-3p are shown. **H.** GSEA analysis showed a same trend as the heatmap, in which pathways down/up-regulated by *Tsc1*-Sh are rescued by miR-495-3p inhibition.

### 2.4. ASO-directed inhibition of miR-495-3p is sufficient to prevent *Tsc1* knockdown induced hyposocial behavior

Next, we assessed whether inhibition of individual miRNAs of the miR-379-410 cluster was able to prevent *Tsc1*-knockdown induced hyposociability. For these experiments, we employed locked nucleic acid (LNA)-modified ASOs (pLNAs), which are efficient miRNA inhibitors and can be easily translated into therapeutic settings, focusing on miR-495-3p (Figure 2), miR-134-5p and miR-485-5p (Narayanan, Levone et al. 2024). We injected female mice in the hippocampus with either Ctrl-Sh or *Tsc1*-Sh, in combination with pLNAs against either the miR-495-3p alone, a combination of miR-134-5p/485-5p, or an unrelated sequence (Ctrl, no known target in mice) three weeks before subjecting them to a behavioral test battery (timeline in Figure 4A). Strikingly, inhibition of miR-495-3p alone was sufficient to completely prevent *Tsc1* knockdown-induced hyposocial behavior in the reciprocal social interaction test (Figure 4B) as well as in the three-chambers social preference test (Figure 4C), whereas hyposocial behavior was preserved in mice treated with the miR-134-5p/miR485-5p pLNA cocktail. Interestingly, while miR-134-5p/miR485-5p inhibition did not change *Tsc1* knockdown-induced impairments in short-term object recognition, miR-495-3p inhibition prevented this phenotype (Figure 4D). No change was observed in either the distance traveled by the pair during social interaction test (Suppl. Figure 6A) or in locomotor activity in the open field test (Suppl. Figure 6B) for any of the pLNA-injected mice. Further, anxiety-like phenotypes, as measured by the time spent in the center of the open field (Suppl. Figure 6C) and the transitions in the light-dark box (Suppl. Figure 6D), were unaltered in these mice.

When validating the successful inhibition of the mature miRNAs by the pLNAs through Taqman RT-qPCRs, we observed that while the expression of miR-495-3p (Suppl. Figure 6E) and miR-485-5p (Suppl. Figure 6F) were strongly reduced by their respective pLNAs, the expression of miR-134-5p was surprisingly slightly increased (Suppl. Figure 6G). The expression of miR-138-5p, used as negative control, was unchanged across the groups (Suppl. Figure 6H).

We next performed poly(A) RNA sequencing on samples from Ctrl-Sh, *Tsc1*-Sh, and *Tsc1*-Sh with miR-495-3p inhibition (*Tsc1*-Sh + pLNA 495). Differential expression analysis revealed widespread transcriptional changes between *Tsc1*-Sh with and without miR-495-3p inhibition. Notably, a substantial subset of genes downregulated by *Tsc1* knockdown was restored to near-control expression levels upon miR-495-3p inhibition. Conversely, genes upregulated following *Tsc1* knockdown were similarly normalized by miR-495-3p inhibition (heatmap, Figure 4E). Among the genes showing this rescue pattern were *Clock* and *Gria2* (Figure 4F), genes correlated with autism (Yang, Matsumoto et al. 2016, Cai, Zhou et al. 2022), and *Pak3* and *Washc4* (Figure 4G), genes correlated to intellectual disorders (Rejeb, Saillour et al. 2008, Courtland, Bradshaw et al. 2021). Interestingly, these genes harbor experimentally validated (Tan, Mu et al. 2017, Capauto, Colantoni et al. 2018, Tan, Tong et al. 2021) or bioinformatically predicted binding sites for miR-495-3p. Interestingly, miR-495-3p binding sites have also been previously implicated in the control of social behavior and autism, pointing towards a broader impact of this miRNA in social behavior outside of the mTORC1 context proposed in this manuscript. We also performed a Gene Set Enrichment Analysis (GSEA, Figure 4H) and observed a similar trend as seen in the heatmap, in which pathways down/up-regulated by *Tsc1*-Sh are rescued by miR-495-3p inhibition.

## 3. Discussion

The observation that social behavior exists on a phenotypic continuum, ranging from the hyposociability of Autism Spectrum Disorder (ASD) to the hypersociability of Williams-Beuren Syndrome (WBS), provides a unique framework for identifying the responsible molecular regulators for this phenomenon. In this study, we identified a novel neurobiological axis where *Tsc1* knockdown in excitatory neurons of the hippocampus, and the consequent mTORC1 hyperactivation, drive social deficits specifically through the upregulation of miR-495-3p, a member of the miR-379-410 cluster. By utilizing acute, shRNA-mediated knockdown of *Tsc1*, we bypassed the confounding effects of severe epilepsy typically seen in constitutive models (Bateup, Johnson et al. 2013). This allowed us to reveal that even modest, localized reductions in *Tsc1* in the hippocampus are sufficient to induce hyposociability and cognitive impairment.

In the mouse liver, *Tsc1* knockdown was recently shown to induce a robust upregulation of multiple members of the miR-379-410 cluster: 18 of the 31 upregulated miRNAs originated from this cluster, with additional changes involving other miRNAs within the same imprinted *Dlk1*-*Dio3* locus (Liko, Rzepiela et al. 2020). The authors attributed this effect to altered DNA methylation at this imprinted region, leading to its transcriptional activation, which is otherwise strongly repressed in the liver, as reflected by the very low baseline expression of these miRNAs. In contrast, neurons express miR-379-410 cluster miRNAs at substantially higher basal levels, which may explain why *Tsc1* knockdown does not produce a comparable global upregulation of the cluster in neuronal tissue. Nevertheless, across multiple cohorts we consistently observe a modest but significant increase in the expression of one cluster member, miR-495-3p. Notably, miR-495-3p has been shown to regulate the Notch/PTEN/AKT signaling (Zhang, Yang et al. 2018), a key pathway implicated in neurodevelopmental disorders, including ASD. Because Notch signaling can indirectly enhance mTORC1 activity by repressing PTEN and promoting AKT activation, changes in miR-495-3p levels may further modulate signaling upstream of mTORC1, potentially contributing to the rescue of *Tsc1* knockdown-associated phenotypes. Interestingly, our RNA sequencing revealed that *Pten*, a known ASD-linked gene (Rademacher and Eickholt 2019), showed a trend towards upregulation in the pLNA-495 injected mice. Moreover, mTORC1 inhibition was shown to rescue Pten deleletion-induced neuronal phenotypes *in vitro* and *in vivo* (D’Amore, Sundberg et al. 2025).

Our findings identify the miR-379-410 cluster as a key molecular effector downstream of mTORC1 in the control of social behavior. In female mice, the hyposocial phenotype induced by *Tsc1* knockdown was fully reversed by pharmacological mTORC1 inhibition with rapamycin and, strikingly, was completely prevented by genetic deletion of the miR-379-410 cluster in excitatory neurons. Conversely, acute pharmacological activation of mTORC1 using NV-5138 phenocopied the *Tsc1* depletion, demonstrating that mTORC1 hyperactivation is sufficient to suppress sociability. Together, these data place the miR-379-410 cluster as a necessary mediator of mTORC1-dependent social behavior deficits. In male mice, *Tsc1* knockdown produced more variable effects, which may reflect a sex-specific baseline regulation of sociability by the miR-379-410 cluster. Indeed, we previously showed that deletion of this cluster alone induces a robust hypersocial phenotype selectively in males (Narayanan, Levone et al. 2024), suggesting a stronger constitutive repressive role in males at baseline. In females, where baseline repression appears less pronounced, perturbations of mTORC1-miR-379-410 signaling may therefore produce more overt behavioral consequences.

Intriguingly, while the miR-379-410 cluster knockout in excitatory neurons rescued social behavior but not memory, the global inhibition of miR-495-3p in all hippocampal cells using Locked Nucleic Acid (pLNA) ASOs mitigated both phenotypes. This suggests that miR-495-3p targets involved in the control of social behavior could be important in non-excitatory cell types, e.g., parvalbumin(Pv)-positive interneurons. In this scenario, Pv-interneurons could interact with *Tsc1*-deficient excitatory neurons to normalize cognitive output.

While previous studies have focused on *Tsc1* in inhibitory interneurons or Purkinje cells, the role of *Tsc1* specifically within hippocampal excitatory circuits remains under-explored. As the hippocampus is vital for social memory and cue processing, a hyperactive mTORC1 pathway may lead to a failure in regulating excitatory synaptic strength, rendering the circuit physiologically incapable of supporting normal social approach. With the *Tsc1*-mTORC1-miR-379-410 axis dysregulated, social deficits may arise from a failure to properly “prune” or regulate excitatory strength, leading to a circuit that is physiologically incapable of supporting normal social approach.

While the current clinical “gold standard” for treating mTOR-related pathologies (such as tumors and epilepsy in Tuberous Sclerosis patients) is rapamycin, its systemic use carries significant metabolic and immunological risks. Our results offer an attractive alternative by providing proof-of-concept for miRNA-based precision therapeutics. Inhibition of the *Tsc1* downstream target miR-495-3p with ASOs potentially “releases the brake” on sociability and rescues cognitive function without disrupting the essential homeostatic functions of the broader mTORC1 pathway. In summary, our study reveals a link between hippocampal *Tsc1* and miR-495-3p, thereby bridging the gap between a major clinically relevant signaling pathway and a localized social regulator. Such intervention could also be relevant in other disorders that include a social aspect, such as schizophrenia and depression, which were also shown to display dysregulated mTORC signaling (Ignacio, Reus et al. 2016, Chadha and Meador-Woodruff 2020). Future studies should focus on validating mRNA as direct targets of miR-495-3p, and use also other *Tsc1*-deficiency models, with the aim of further clarifying the pathophysiology of ASD and refining the potential for emerging miRNA-based interventions.

## 4. Methods

### 4.1. *In vivo* studies

#### 4.1.1. Ethical statement

All animal experiments were conducted in strict accordance with the Swiss Federal Act on Animal Protection and the corresponding ordinances. Experimental procedures were reviewed and approved by the Cantonal Veterinary Office of Zurich under licenses ZH194/2021 and ZH093/2025.

All efforts were made to comply with the principles of the 3Rs (Replacement, Reduction, and Refinement). Animal use was minimized through appropriate experimental design and statistical planning. All procedures were refined to minimize animal distress and ensure the highest standards of animal welfare.

#### 4.1.2. Animals

Mice were housed in groups of 2-5 per cage, with food and water *ad libitum*, and kept in an animal room with inverted light-dark cycle (lights off at 8:00, on at 20:00), and all tests were performed during their dark phase. Males and females C57Bl/6 mice were used, and according to the specific experiment, different genotypes were used (described in the specific results section and figure legends): wild types (WT, purchased from Janvier, France) or Emx1-Cre-driven conditional knockout of the miR-379-410 cluster (cKO, and their control/CTL, bred in-house). Experimental mice were tested at postnatal week (PNW) 12-20, according to each experiment. CTL/cKO mice were genotyped in-house using the KAPA mouse Genotyping kit (KAPA Biosystems). Mice were handled daily for 5 min for a week before the behavioral experiments began. During behavioral experiments, mice were housed individually and allowed to acclimatize to the experimental room in a holding cage for 30 min (or 1 h for the social behavior tests) before the beginning of the task. At the end of each experiment, mice were placed back into their original home cage with their littermates. 10 mL/L detergent (Dr. Schnell AG) was used to clean the equipment in between trials. Behavioral testing were performed from the least to the most stressful and experimenters were blinded during the analyses of behavioral videos that required manual scoring.

#### 4.1.3. Stereotaxic surgeries

After quick anesthesia induction by 5% isoflurane inhalation in oxygen (1 L/min), mice were transferred to a stereotactic frame, having their ears fixed using ear bars, while laying on a warmed plate and wearing a mouth/nose mask with a flow of 1.5-2.5% isoflurane in 200 mL/min oxygen. EMLA cream was used as local anesthetic in the ears prior to placing the ear bars. Meloxicam (5 mg/kg in saline) was subcutaneously injected, and a Vitamin A cream was applied to protect eyes from drying. After the mouse head was shaved and cleaned with Betadine, an incision was made. Following the bregma and lambda coordinates, bilateral injections were performed in the dorsal/intermediate hippocampus at the coordinates (from bregma): AP -2.06 mm, ML ±1.53 mm and DV -1.51 mm.

Plasmids were designed and purchased from VectorBuilder, and viral vectors (rAAV-PHP.eB) were produced by the ETH Vector and Virus Production Platform (see Suppl. Table 1 for more details). Mice were injected with 1 µL of virus per hemisphere using a fine glass capillary over 5 min, followed by an additional 1 min to allow viral diffusion. The capillary was then withdrawn slowly over 3 min. In a subset of experiments, mice received pLNAs in combination with the virus (see Suppl. Table 2 for more details). For these injections, pLNAs were diluted to 300 µM in PBS and mixed with the viral preparation immediately before injection (1 µL virus + 0.5 µL pLNA), with a total volume of 1.5 µL being injected per hemisphere.

Local analgesia was provided by suturing the skin while adding Lidocain and Bupivacain (2 mg/kg each) drops on the wound site. Mice were then allowed to recover from the surgery before being group housed again. A second subcutaneous injection of meloxicam (5 mg/kg in saline) was performed 8-12 h post-surgery, and animals had paracetamol added to their drinking water for the next 48 h. Daily postoperative health checks were carried out daily over the 3 following days and weekly after that.

#### 4.1.4. Drug injections

The mTORC1 activator NV-5138 and inhibitor Rapamycin were diluted in DMSO at the concentrations of 100 µM and 10 µM, respectively, aliquoted and frozen at -80°C. On the day of the injection, aliquots were thawed and diluted in saline (for a final saline concentration of 0.9%). Control mice were injected with 0.9% saline and the same amount of DMSO as the respective drug. For each drug or control, each mouse received 0.20 – 0.3 mL of the final solution, depending on their body weight. In total, treated mice received 12 mg/kg of the NV-5138 or 5 mg/kg of Rapamycin. After the injection, mice were group housed again for 2 h and then singly housed 1 h prior to being subjected to the reciprocal social interaction test.

#### 4.1.5. Behavioral testing

##### 4.1.5.1. Open field

Mice were placed in an open-field arena (45 × 45 × 40 cm; TSE Systems, Bad Homburg, Germany) and allowed to explore for 10 min. Sessions were videorecorded, and locomotor activity and time spent in the center were automatically quantified using TSE VideoMot2 software (TSE Systems).

##### 4.1.5.2. Reciprocal social interaction test

Mice were single-housed for 1 h before testing. Two unfamiliar mice of the same genotype (and treatment, when applicable) were placed together in the open-field arena and allowed to interact for 10 min. After a further 1 h of single housing, animals were tested again with a different unfamiliar mouse of the same genotype or treatment. Interactions were videorecorded, and time spent in social behaviors (nose and ano-genital sniffing, scratching, seeking/following) was scored manually. No aggressive behavior was observed during sessions.

##### 4.1.5.3. Three chambers test

Social preference and social memory were assessed using a three-chambers apparatus (60 × 43 × 22 cm; 20 cm per chamber). The test consisted of three 10-min sessions separated by 30-min intervals. During habituation, mice freely explored the arena with an empty wire cage in each side chamber. In the social preference session, one cage contained an unfamiliar sex- and age-matched mouse, while the other contained an object (lego tower). In the social memory session, mice were exposed to the previously encountered mouse and a novel unfamiliar mouse. Sessions were videorecorded, and time spent sniffing/directly interacting with each cage during the second and third sessions was scored manually. Social preference was expressed as time spent sniffing the social cage divided by the empty cage; social memory as time spent sniffing the novel mouse divided by the familiar mouse.

##### 4.1.5.4. Light-dark box

Mice were placed in a light-dark box (45 × 30 × 25 cm) consisting of a light compartment (28.5 cm) and a dark compartment (16.5 cm) connected by a small door and allowed to explore for 10 min. Sessions were videorecorded, and the number of transitions between compartments was automatically quantified using TSE VideoMot2 software (TSE Systems)

##### 4.1.5.5. Novel object recognition test

Mice were habituated to the testing box on the day prior to the experiment. On the day of the experiment, mice were first exposed to two identical objects in an open-field arena (45 × 45 × 40 cm) for 5 min, followed by a 15-min intertrial interval. In the test session, mice were reintroduced to the arena containing one familiar and one novel object for 5 min. Objects were selected and randomized according to a published protocol (Leger, Quiedeville et al. 2013) and validated in a separate cohort to exclude innate object preference. Exploration time was scored manually and expressed as time spent sniffing the novel object divided by the familiar object.

##### 4.1.5.6. Fear conditioning test

Fear conditioning was performed in TSE conditioning chambers (30 × 30 × 25 cm; Plexiglas walls, metal grid floor). During training, mice were placed in the chamber for 3 min and then received a single foot shock (2 s, 0.8 mA) with a cue (beep), after which they were returned to their home cage. Twenty-four hours later, mice were re-exposed to the same context with the cue for 5 min, and freezing behavior was automatically quantified using TSE VideoMot2 software (TSE Systems).

#### 4.1.6. Mouse tissue collection

After the end of behavioral experiments, mice were euthanized and tissue was collected for molecular analyses. For RNA extractions, mice were quickly anesthetized with 5% isofluorane, decapitated, the hippocampi were dissected under a fluorescent microscope (to enrich for GFP-positive area) and snap-frozen in liquid nitrogen and kept at -80°C (see next steps in item 2.3.1 below). For protein assays, hippocampi were dissected as above and collected in RIPA (for bulk assay) or homogenization buffer (for crude synaptosomal extractions) (see next steps in item 2.3.5 below). For immunohistochemistry, mice were perfused. Briefly, mice were anesthetized, perfused with ice-cold PBS for 5 min and then with ice-cold 4% PFA (pH 7.4) for 5 min. Brains were dissected and post-fixed in 4% PFA overnight. For three days, brains were transferred to sucrose solution at 4°C (in PBS), increasing in concentration: 10, 20 and 30%. After sinking in the 30% sucrose solution, brains were frozen at -80°C embedded in Optimal Cutting Temperature (O.C.T.) (see next steps in item 2.3.4 below).

### 4.2. *In vitro* studies

#### 4.2.1. Neuro2a cultures and transfections

Neuro2a cells were cultured in DMEM high glucose culture medium supplemented with 10% bovine serum albumin and 1% antibiotics (Penicillin/Streptomycin). On the day before the transfection, cells were trypsinized and 300,000 cells were plated per well on 6-wells plates. The next day, medium was changed to an antibiotic-free DMEM (1.5 mL). A mix between OptiMEM (500 µL per well), Lipofectamine 2000 (Thermo Fisher, 7.5 µL per well), and plasmid DNA (2.5 µg per well) was vortexed and 0.5 mL was added per well dropwise 20 min later. On the next day, cells were washed, trypsinized, and re-plated at 200,000 cells per well in 6-wells plates. Two days later, cells were washed once in PBS and then collected in Trizol for RNA extractions or fixed for 15 min in 4% PFA / 4% sucrose (Merk / SigmaAldrich) for immunocytochemistry. Plasmids used for transfection are listed in Suppl. Table S1.

### 4.3. Molecular Analyses

#### 4.3.1. RNA extraction

Frozen hippocampi samples were thawed, 1 mL TRIzol Reagent (Thermo Fisher) was added, and tissue was homogenized at 4°C in a tissue lyser bead mill (Qiagen) for 2 min at 20 Hz. Subsequently, 200 µL of chloroform were added and homogenate was incubated for 3 min at room temperature (RT), before being centrifuged at 12,000 × g for 15 min at 4°C. Aqueous phase was transferred to a fresh tube. A second clearing was performed by adding the same amount of chloroform to the aqueous phase and centrifugating at 12,000 × g for 15 min at 4°C. The cleared aqueous phase was then mixed with 2 µL of Glycogen to allow RNA visualization (Ambion) and 500 µL of 100% isopropyl alcohol, incubated at RT for 10 min and then centrifugated. RNA pellet was washed twice with 1 mL of 75% ethanol (5 min centrifugations at 7,500 × g at 4°C in between washes). After 15 min drying at RT, pellets were dissolved in 20 µL of nuclease-free H_2_O (Ambion) by pipetting up and down. To ensure complete resuspension of the pellet, samples were incubated for 10 min at 56°C in a heat block. RNA purity and quantity were determined with a Nanodrop 1000 spectrophotometer (DeNovix, Life Science Technologies).

#### 4.3.2. RT-qPCR

Isolated RNA was treated with the TURBO DNase enzyme (Thermo Fisher) according to manufacturer’s directions. For detection of mRNAs, DNAse-treated RNA samples were reverse-transcribed using the iScript cDNA synthesis kit (Bio-Rad), according to the manufacturer’s recommendations, on a C1000 TouchTM Thermal Cycler (BioRad). Quantitative real-time PCR was performed on a CFX384 Real-Time System (BioRad), using iTaq Universal SYBR Green Supermix (Bio-Rad) according to manufacturer’s instructions. For the detection of miRNAs, DNAse-treated RNA samples were reverse-transcribed using the Taqman MicroRNA Reverse Transcription Kit (Thermo Fisher Scientific). Quantitative real-time PCR was performed using Taqman Universal PCR Master Mix (Thermo Fisher Scientific), according to manufacturer’s instructions. Primers were obtained from Microsynth (mRNA) or Thermo Fisher (miRNA) and their sequences are listed in Suppl. Table 3. Each sample was measured in triplicate. Using the mean Ct-value of these triplicates, real-time RT-qPCR data were analyzed by normalizing each gene of interest to the housekeeping gene U6 (for miRNAs and mRNAs). The relative RNA expression was calculated as follows: = POWER (2-(Ct of gene - Ct of housekeeping)). After that, values were normalized by the control group (which was set at 100%).

#### 4.3.3. Immunocytochemistry

After fixation, Neuro2a cells were three times in PBS and then blocked in blocking buffer (10% normal goat serum [NGS] and 0.3% Triton in PBS) for 1 h at room temperature. Cells were then incubated with primary antibodies (TSC1) diluted in antibody buffer (5% NGS and 0.3% Triton in PBS) for 1 h at room temperature. After 3 washes with 0.1% Triton/PBS (PBS-T), cells were incubated with secondary antibodies (Alexa Fluor 647, diluted in antibody buffer) for 1 h at room temperature. After 3 washes with PBS-T, cells were incubated with Hoechst 33342 (diluted in PBS) for 10 min at room temperature. Cells were then washed 3 times with PBS, once with dH_2_O, and coverslips were mounted onto microscope slides using Aqua-Poly/Mount (Chemie Brunschwig) (see Suppl. Table 4 for more information on antibodies and dilutions).

#### 4.3.4. Immunohistochemistry

Frozen brains (-80°C) were sliced at 50 mm using a cryostat and slices were kept in anti-freeze solution (25% PBS 0.2M, 30% ethylene glycol, 25% glycerol and 20% dH_2_O). Free-floating brain slices were washed twice in PBS for 5 min and then incubated in blocking buffer (10% NGS and 0.3% Triton in PBS) for 1 h at room temperature. Slices were then transferred to a primary antibody solution (TSC1, diluted in antibody buffer) for 1 h at room temperature. Slices were washed three times in PBS-T and then incubated in a secondary antibody solution (Alexa Fluor 647, diluted in antibody buffer) for 1 h at room temperature. Slices were washed three times in PBS-T and then counterstained with Hoechst 33342 (diluted in PBS) for 10 min. Slices were finally washed three times in PBS and then mounted onto charged slides and a glass coverslip was mounted using Aqua-Poly/Mount (see Suppl. Table 4 for more information on antibodies and dilutions).

#### 4.3.5. Protein assays

##### 4.3.5.1. Bulk protein extracts

Proteins from bulk hippocampal tissue were lysed using RIPA buffer (50 mM Tris-HCl pH 7.5, 150 mM NaCl, 1% NP-40, 0.5% sodium deoxycholate, and 0.1% SDS) containing proteinases and phosphatases inhibitor. Tissue was mashed using a pestle and left on ice for 30 min before being centrifuged at 16,000 g for 10 min at 4°C to remove unsoluble proteins and cell debris. Supernatant was transferred to a new tube and protein concentrations were determined using a bicinchoninic acid assay (BCA). Samples were then diluted to 0.5-1 mg/mL with nuclease-free water and 4x Laemmli Sample Buffer (Bio-Rad) containing 10% β-mercaptoethanol. Proteins were then denatured by boiling at 95°C for 5 min in a thermoblock.

##### 4.3.5.2. Preparation of crude synaptosomal extracts

Hippocampi of mice were dissected and washed in ice-cold PBS (hippocampi of 2 mice were pooled per sample). Tissue was homogenized in 500 µL of homogenization buffer (0.35 M sucrose, 0.5 mM EGTA, 10 mM Tris-HCl, protease inhibitor cocktail and phosphatase inhibitor) with a teflon-coated Dounce-Potter homogenizer by eight up and down strokes. Homogenate was transferred to 1.5 mL tubes and centrifuged at 2,000 g for 5 min at 4°C to discard nuclei and cell debris (pellet). Supernatant was transferred to a new tube and centrifuged at 30,000 × g for 10 min at 4°C to pellet a crude synaptosome-containing fraction. Pellets were resuspended in RIPA buffer containing proteinases and phosphatases inhibitors.

Protein concentration was measured by BCA and samples were diluted to 0.5-1 mg/mL with nuclease-free water and 4x Laemmli Sample Buffer (Bio-Rad) containing 10% β-mercaptoethanol. Proteins were then denatured by boiling at 95°C for 5 min in a thermoblock.

##### 4.3.5.3. Western blot

Proteins (10 µg) were separated on a 10% polyacrylamide gel in an SDS-PAGE running buffer (25 mM Tris, 192 mM glycine, 0.1% SDS). After electrophoresis, proteins were transferred to a nitrocellulose membrane using a Trans-Blot Turbo system (Bio-Rad), according to manufacturer’s protocol. Membranes were blocked in 5% non-fat dry milk (for non-phosphorylated proteins) or 4% bovine serum albumin (for phospho-S6), diluted in Tris-buffered saline solution (50 mM Tris, 150 mM NaCl, pH 7.5) containing 0.1% Tween 20 (TBS-T) and then incubated in primary antibody solution diluted in blocking buffer overnight at 4°C. Membranes were washed 4 times TBS-T before incubating them in HRP (horseradish peroxidase)-conjugated secondary antibody solution at room temperature for 1 h (see Suppl. Table 4 for more information on antibodies and dilutions). Following incubation, membranes were washed 4 times in TBS-T, then developed with the ClarityTM Western ECL Substrate (Bio-Rad) according to manufacturer’s guidelines and visualized with the ChemiDocTM MP, Imaging System (BioRad). Subsequent quantification of the protein levels was performed using the software Image Lab 6, and the intensity of the protein of interest was normalized by the intensity of Tubulin.

#### 4.3.6. Image acquisition and analysis

Immunocyto- and histochemistry images were acquired with a confocal laser-scanning microscope (CLSM 880, Zeiss) using z stack (set interval 0.45 mm) with a 63x oil objective at a variable resolutions, depending on experiment (Figure 2A: 4576 × 7576; Figure 2B: 1228 × 1228; Suppl. Figure 1B: 1128 × 1128). The maximum intensity projection of the z stack images was used. Fluorescence intensity was measured in at least 3 images per animal in three animals (or per coverslip area, done in triplicates for Neuro2a). Results were normalized by the average intensity of the control.

#### 4.3.7. Poly-A RNA sequencing analysis and Geneset Enrichment Analysis (GSEA)

Reads were mapped to the GRCm39 genome and genes were quantified (ensembl release 112) with STAR 2.7.11a. Genes were filtered with edgeR::filterByExpr using a minimum count of 20, and differential analysis was performed with edgeR 4.6.3. As characteristic patterns of choroid plexus contamination was observed in some samples, we computed a contamination score based on the average logCPM expression of the genes *Ttr*, *Igfbp2*, *Prlr*, *Enpp2*, *Sostdc1*, *Ecrg4*, *Kl*, *Clic6*, *Kcne2*, *F5*, *Slc4a5*, *Aqp1*, and was included as a covariate in the differential expression model.

Gene-set enrichment analysis (GSEA) was performed using multi-level procedure from the fgsea package version 1.34.2, with a minimum size of 5. The synGO release from 20231201 was used. For the molecular signature database, the mouse collections CP::KEGG_MEDICUS, GO:BP, GO:CC, and GO:MF were used from the msigdbr package version 25.1.0.

### 4.4. Data visualization and Statistical analysis

All data were analyzed and plotted using GraphPad Prism 10 for Windows (GraphPad Software, San Diego, California USA, www.graphpad.com). For two-groups comparison, data were analyzed using unpaired two-tailed Student’s t-test. For three-groups comparison, data were analyzed using one-way ANOVA followed by Tukey post-hoc test, when appropriate. For four-groups comparison, either one- or two-way ANOVA (depending on the experimental design) followed by Tukey post-hoc test (when appropriate) was used. Statistical results can be found in respective Figure legends and are summarized in Suppl. Tables 5 and 6. Figure legends also display the sample size, which represents the number of biological samples per group; if groups have different size, the single sample number per group is shown in the order they appear in the graph, prior to the p value. Data are presented as mean ± SD. p < 0.05 was considered statistically significant (in graphs, n.s. p > 0.05, *p < 0.05, **p < 0.01 and ***p < 0.001).

## Acknowledgements

We thank Dr. Roberto Fiore and Alessandra Lo Bianco for technical advise and Cristina Furler for laboratory support. This work was funded by a grant from the Swiss National Science Foundation (SNSF), grant no. 310030_205064.

## Author Contributions

BL planned and performed experiments and wrote the paper; NS and PD performed experiments; PLG performed RNA sequencing analyses; GS supervised the project and revised the paper.

## Data availability

RNA sequencing data is being deposited to Gene Expression Omnibus (GEO). Accession number and password available upon request.

